# Electronic cigarettes induce mitochondrial DNA damage and trigger toll-like receptor 9-mediated atherosclerosis

**DOI:** 10.1101/2020.08.15.252494

**Authors:** Jieliang Li, Do Luong Huynh, Moon-Shong Tang, Hannah Simborio, Jing Huang, Beata Kosmider, Michael B. Steinberg, Le Thu Thi Le, Kien Pham, Chen Liu, He Wang

## Abstract

**Objective:** Both electronic cigarette (e-cig) use and toll-like receptor 9 (TLR9) activation have been implicated in promoting atherosclerosis. In this study we aimed to investigate the causative relationship of e-cig exposure on TLR9 activation and atherosclerosis development.

**Approach and Results:** Eight-week-old ApoE-/- mice fed normal chow diet were exposed to e-cig vapor (ECV) for 2 h/day, 5 days/week for 16 weeks. We found that ECV exposure significantly induced atherosclerotic lesions as examined by Oil Red O staining of aortic root and greatly upregulated TLR9 expression in classical monocytes and in the atherosclerotic plaques, which the latter was corroborated by upregulated TLR9 expression in human femoral artery atherosclerotic plaques in e-cig smokers. Intriguingly, we found a significant increase of damaged mitochondria DNA level in the circulating blood of ECV exposed mice. Furthermore, administration of TLR9 antagonist prior to ECV exposure not only alleviated atherosclerotic lesion and the upregulation of TLR9 in plaques, but also attenuated the increase of plasma levels of inflammatory cytokines, reduced the accumulation of lipid and macrophages, and decreased the frequency of blood CCR2+ classical monocytes. Surprisingly, we found that the cytoplasmic mtDNA isolated from ECV extract-treated cells can greatly enhance the expression of TLR9 in reporter cells.

**Conclusion:** E-cig induces mtDNA damage and the mtDNA in circulating blood stimulates the expression of TLR9, which elevate the expression of proinflammatory cytokines in monocyte/macrophage and consequently lead to atherosclerosis. Our results raise the possibility that intervention of TLR9 activation is a potential pharmacologic target of ECV-related inflammation and cardiovascular diseases.

## Introduction

Atherosclerosis is a disease of large- and medium-sized arteries that is characterized by the formation of subendothelial plaques consisting of necrotic cores, accumulated modified lipids, extracellular matrix, smooth muscle cells (SMCs), endothelial cells (ECs), leukocytes, and foam cells.^1^ Atherosclerosis is the pathological basis of a variety of cardiovascular diseases (CVDs), which are the leading causes of death worldwide.^2^ Cigarette smoking is a well-established major risk factor for CVDs, recognized as such for decades.^3^ A large body of evidence has indicated that smoking accelerates atherosclerosis and induces a hypercoagulable state.^4–6^ In the past 10 years, electronic cigarette (e-cig) use has increased rapidly in the United States^7, 8^ whereas the impact of e-cigs on CVDs has been reported only just recently. A randomized, double-blinded pilot study revealed that e-cigs worsen peripheral and central hemodynamics as well as arterial stiffness.^9^ Another study reported that e-cig aerosol increases arterial stiffness and oxidative stress, although to a lesser extent than a single conventional cigarette.^10^ Studies in animals have found that e-cigs have both rapid and long-term negative effects on arteries, inducing cardiac dysfunction and atherosclerosis.^11, 12^ However, despite the reported association of e-cigs with atherosclerosis, the underlying mechanisms of the pro-atherosclerotic effects of e-cigs remain unclear.

Toll-like receptors (TLRs) are a class of essential components of the innate immune system that recognize distinct kinds of pathogen-associated molecular patterns (PAMPs). Activation of TLRs triggers an intracellular signaling cascade mediated through MyD88 or TRIF, leading to the production of pro- and anti-inflammatory cytokines.^13^ TLR9 activation of murine and human macrophages has been shown to stimulate their inflammatory responses (TNF-α, IL-6) and accelerate their transformation into foam cells, an indication of plaque build-up.^14, 15^ Activation of TLR9 by relatively high dosage of oligonucleotide ODN1826 was found to increase plaque formation^16^ and TLR9 inhibition alleviated atherosclerotic progression in ApoE^−/−^ mice fed a high-fat diet.^17^ Recently, evidence has accumulated to indicate that TLR9 plays a pivotal role in the development of vascular inflammation through proinflammatory activation of macrophages in angiotensin II-induced atherosclerosis.^18^ In addition, pharmacological blockade of TLR9 can attenuate atherosclerotic lesion formation, suggesting TLR9 may serve as a potential therapeutic target for atherosclerosis.^18^

TLR9 also recognizes endogenous self-DNA termed damage-associated molecular patterns (DAMPS) from injured mammalian cells.^19^ For example, mitochondrial DNA (mtDNA) is able to activate TLR9 and induces podocyte apoptosis.^20^ TLR9 activation by circulating mtDNA is reported to contribute to elevated arterial pressure and vascular dysfunction in spontaneously hypertensive rats (SHR).^21^ In fact, direct DNA damage is a critical mechanism underlying smoking-associated tumorigenesis (e.g. lung cancers).^22–25^ Smoke exposure is associated with increased mtDNA damage both *in vitro* and *in vivo*.^26, 27^ MtDNA damage is implicated in promoting adult atherogenesis by prenatal environmental tobacco smoke exposure in ApoE^−/−^ mice.^28^ The detrimental effects of ecigs on genomic DNA have also been recently identified. E-cig vapor (ECV)-exposed cells have significantly increased DNA strand breaks along with increased rates of apoptosis and necrosis.^29^ E-cig aerosols can suppress cellular antioxidant defenses and induce significant oxidative DNA damage.^30^ ECV damages DNA and reduces DNA-repair activity in mouse organs as well as in human cells.^31^ However, effect of e-cig on mtDNA damage, and the biological sensor of the released mtDNA remains elusive. Importantly, no current study establishes the causal relationship between mtDNA damage-TLR9 activation during atherosclerotic process even in the setting of traditional cigarette smoking, let alone e-cig vaporing. In the present study, we examined the impact of ECV on DNA damage and the activation of TLR9 in ApoE^−/−^ mice and investigated whether TLR9 blockade could alleviate e-cig induced development of atherosclerosis.

## Materials and Methods

### Animals

Eight-week-old mice (n = 5~10 per group) were subjected to whole-body exposure. The control mice were exposed to HEPA filtered room air. E-cig aerosol smoke was generated from Platinum V2 RED E-liquid (classic tobacco flavor containing 2.4% nicotine) by a Teague smoking machine (Model TE-2, Teague Enterprises, Woodland, CA). The aerosol was generated by heating 2 mL e-liquid as we previously described.^32^ Mice received 2 h exposures per day for 5 successive days per week, over a 16 week period. For the animal group with pharmacological TLR9 blockade, mice received intraperitoneal injections of TLR9 antagonist IRS869^33^ (5 mg/kg, 5’-TCCTGGAGGGGTTGT-3’; IDT, Coralville, IA) twice per week and 1 h prior to e-cig exposure. The mice were euthanized with pentobarbital sodium (200 mg/kg) by intraperitoneal injection after the last exposure. Blood and tissues were collected and stored for further assays. Animal handling and experimentation were in accordance with the recommendation of the current NIH guidelines and were approved by the Temple University School of Medicine Institutional Animal Care and Use Committee.

### Flow cytometry

Whole blood cells were stained and analyzed on a FACSCanto LSRII flow cytometer (BD Biosciences). Isotype controls were used to set appropriate gates. Detailed staining protocol and antibodies are described in supplementary materials. Data were analyzed acquired with FACSDiva (BD Biosciences) and analyzed with FlowJo 6.4.7v10.6 (Tree Star, Ashland, OR). For all samples, approximately 20,000 cells were analyzed to generate scatter plots. All events were collected from each sample to generate the scatter plots.

### Multiplex ELISA for mouse cytokines

Aliquots of plasma from mice were analyzed using mouse cytokine 9-Plex ELISA kit from Boster Biological Technology (Pleasanton, CA) according to manufacturer’s instructions.

### Histological studies

The formalin fixed heart and whole aorta were dehydrated in ethanol and embedded in paraffin, and 5-μm serial sections were stained with hematoxylin and eosin (Millipore-Sigma, Burlington, MA). Proximal aortic sections were obtained within the root. Sections were evaluated blindly to score for pathological changes. The expressions of TLR9, adhesion molecule VCAM-1 and F4/80 were examined by immunohistological staining with individual antibodies using routine procedure. Experiments were repeated three or more times, with 4-5 animals per group.

### Atherosclerotic lesions

Another subset of hearts and proximal aortas were dehydrated in 30% sucrose at 4°C overnight and embedded in OCT compound (Tissue-Tek, Elkhart, IN), frozen on dry ice, and then stored at −80°C until sectioning. Serial 10-μm-thick cryosections (50~60 sections per mouse) were cut through the ventricle from the appearance to the disappearance of the aortic valves. Every other section was collected on poly-D-lysine-coated slides. The deposition of lipids was determined with Oil Red O (ORO) staining (Sigma-Aldrich, St. Louis, MO) and counterstained with hematoxylin and fast green for visualization of atherosclerotic lesions. Atherosclerotic lesions were quantified by taking an average of three cross sections from each aorta spaced 50 μm apart using Image Pro plus 6.0 software (Media Cybernetics, Rockville, MD). Briefly, the lesions were circled, and the area of each lesion for the quantified section on the slide was exported to a spreadsheet (in μm^2^). This was repeated for each of the slides, and the sum of the lesions area was calculated (μm^2^/section) for each mouse.

### Measurement of cytoplasmic DNA lesion and plasma DNA damage

Murine monocyte/macrophage RAW 246.7 cells, maintained in DMEM (Thermo Scientific) plus 10% FBS and 1% antibiotics, were seeded in 60 mm cell-culture dish at density 2 x 10^6^ cell/dish. After 24 h incubation, cells were treated with or without ECV extract (0.5~2.5% v/v; equal to 2.5~12.5 cigarettes/day for conventional cigarettes normalized by nicotine intake which corresponds to the realistic situation of human daily exposure to cigarettes^34^) for 48 h and cytoplasmic DNA was extracted according to the protocol as previously described.^35^ The cytoplasmic DNA lesion and plasma levels of DNA damage were measured using the DNA Damage Competitive ELISA (Thermo Scientific) according to the manufacturer’s instruction. Both plasma DNA damage and cytoplasmic DNA lesion were expressed as the concentration of 8-OHdG.

### Measurement of plasma mtDNA/nDNA ratio and cytoplasmic mtDNA

For measurement of *in vivo* plasma mtDNA/nDNA ratio, mice plasma was first subjected to cell-free DNA (cfDNA) extraction using Apostle minimax cfDNA extraction kit (ApostleBio, CA, US). Both *in vitro* cytoplasmic DNA and in *vivo* plasma cfDNAs were then used as templates for SYBR qPCR (Applied Biosystem, CA) with primers (Supplementary Table S1). The plasma mtDNA/nDNA ratio was calculated as the ratio of mtCO-1/18S rDNA. The relative fold-change of cytoplasmic mtDNA by ECV extract in macrophages was calculated using the 2^-ΔΔCt^ method as compared to control-treated cells.

### *In vitro* analysis of TLR9 activation

To evaluate the TLR9 activation potency of the cytoplasmic DNA, we used engineered HEK293 cell lines (HEK-Blue) that stably express either mouse TLR9 (mTLR9) and an NF-κB reporter gene (Invivogen, San Diego, CA). In brief, HEK mTLR9 blue cells were maintained in selection medium (DMEM complete plus with Blasticidin 30 μg/mL and Zeocin 100 μg/mL) and 100 μL of cells in HEK blue detection medium were seeded in 96 well-plate (at density 2 x 10^4^ cells/ well), followed by adding 20 μL of water or mtDNA (final treatment concentration 25 ng/μL). After 18-24 h incubation, the absorbance was measured at 620-655 nm.

### TLR9 expression in human femoral artery atherosclerotic plaques

Femoral artery atherosclerotic lesions were identified from non-smokers and e-cig smokers (>6 months). The expression of TLR9 was examined by immunohistological staining using routine procedure. Informed consent was obtained from all subjects. The procedures used in this study conform to the principles outlined in the Declaration of Helsinki.

### Data analysis

All data were collated using Microsoft Excel Software and analyzed using GraphPad Prism 5.0 (GraphPad, La Jolla, CA). All data are represented as the mean values with error bars representing standard error of the mean (SEM). Samples were compared using the unpaired Student t test (two-tailed). Differences were considered significant when P<0.05.

## Results

### ECV exposure increased atherosclerosis development and TLR9 expression

We first compared atherosclerotic lesion progression in ECV-exposed ApoE^−/−^ mice with air-exposed mice after 16 weeks. As shown in **Fig. 1A**, the results of en face ORO staining demonstrated a significant increase of atherosclerosis burden in ECV-exposed ApoE^−/−^ mice (P<0.001). ECV exposure also increased the intimal microscopic lesions in the cross-sectional aortic roots as evidenced by HE staining in **Fig. 1B** (P<0.001). Intriguingly, we observed that there was significantly higher expression of TLR9 in the total plaque area in the atherosclerotic lesions in ECV-exposed mice than air-treated mice (**Fig. 1C**). More importantly, in the femoral arterial plaques from e-cig smokers (>6 months), there was higher TLR9 expression as compared to that in non-smoking-related plaques in non-smokers (**Fig. 1D**). These results indicate an undetermined role of TLR9 played in e-cig smoking associated atherosclerosis.

**Figure 1.**
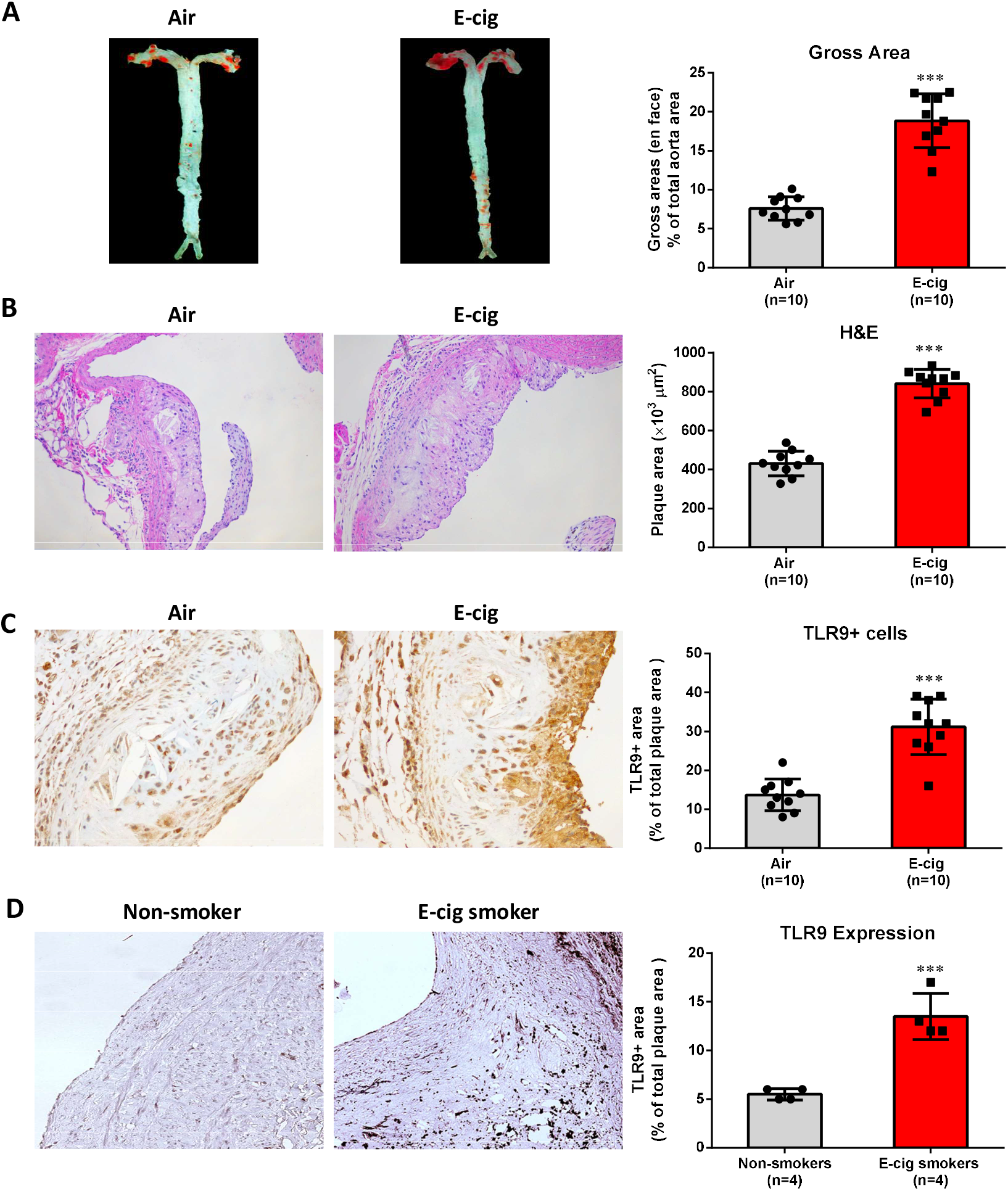
E-cig vapor exposure increased TLR9 expression in atherosclerotic plaques in normal chow-fed ApoE^−/−^ mice, substantiated by the upregulation of TLR9 expression in femoral arterial plaques in e-cig smokers as compared to that in non-smoking-related plaques in nonsmokers. (A) Representative pictures of en face Oil Red O (ORO) staining of freshly dissected intact aorta and quantification analysis of atherosclerosis burden. 8-week-old mice were exposed to air or e-cig vapor for 2 h a day, 5 days per week for 16 weeks. (B) Hematoxylin-eosin (H&E) staining to measure the intimal microscopic lesions in the cross-sectional aortic roots. (C) TLR9 positive cells in atherosclerotic plaques in ApoE^−/−^ mice. Cross-sectioned aortic roots were subjected to TLR9 immunohistochemistry staining. Data are shown as mean ± SEM and statistical analysis was performed using the Students’ t-test. ***P < 0.001. (D) TLR9 expression in the femoral arterial plaques from e-cig smokers (>6 months) as compared to in non-smoking-derived plaques in non-smokers. Data are shown as mean ± SEM (n=4 per group) and statistical analysis was performed using the Students’ t-test. ***P < 0.001.

### TLR9 blockade attenuated ECV-exacerbated atherosclerosis in ApoE^−/−^ mice

We next sought to investigate whether TLR9 blockade can inhibit ECV-induced atherosclerosis development. Mice in the ECV group were administrated with 5 mg/kg of IRS869 prior to ECV exposure. It was found that ECV significantly increased the lipid deposition (~2-fold; **Fig. 2A**) in the atherosclerotic lesion area and enhanced the intimal lesions (~1.9-fold) (**Fig. 2B**) in ApoE^−/−^ mice after 16-week exposure as compared to control mice. In contrast, administration of mice with TLR9 antagonist IRS869 prior to ECV exposure significantly attenuated ECV-exacerbated atherosclerosis development (lipid deposition reduced by 50%; and intimal microscopic lesion decreased by 23%; all P<0.05) (**Fig. 2A, B**).

**Figure 2.**
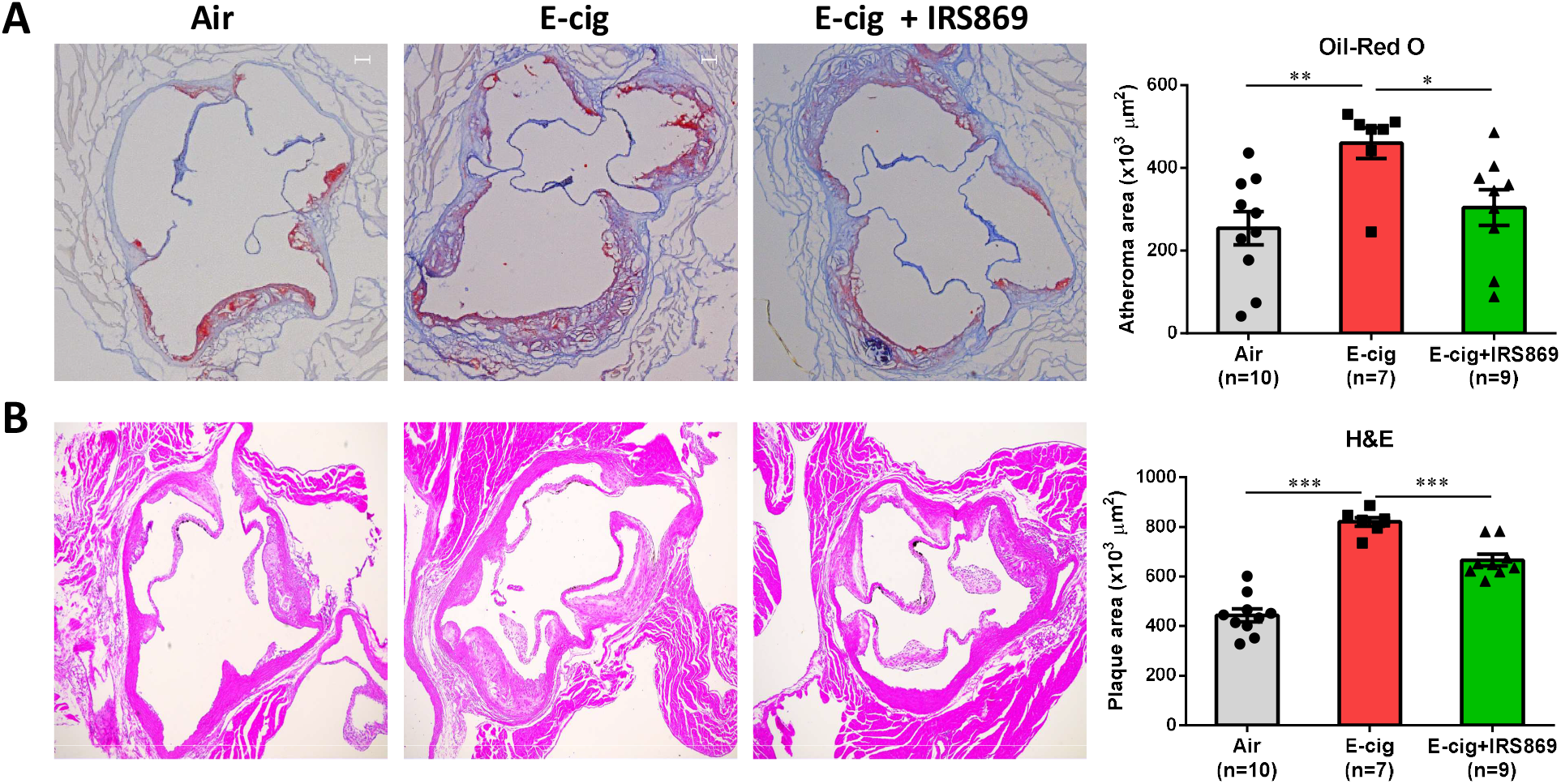
TLR9 inhibitor attenuated ECV-exacerbated atherosclerotic lesion development TLR9 inhibitor in ApoE^−/−^ mice. (A) Oil Red O (ORO) staining of the cross-sectioned aortic roots. Lipid deposition in plaques was quantified as ORO-positive lesion areas and expressed as percentage to the total surface area of the atheroma. (B). Cross-sectional hematoxylin-eosin staining to measure the intimal microscopic lesions in the aortic roots. N means the number of mice in each group. Data are shown as mean ± SEM and statistical analysis was performed using the Students’ t-test. *P < 0.05; **P < 0.01; ***P < 0.001.

### ECV enhanced the expression of TLR9 in classical monocytes, which could be reduced by TLR9 inhibition

Smoking has been shown to activate human monocytes, leading to increased adhesion to the endothelium.^36^ We next determined whether monocyte activation was involved in e-cig-mediated atherosclerotic development. As shown in **Fig. 3A**, ECV significantly increased the percentages of classical monocytes and nonclassical monocytes in the blood of ApoE^−/−^ mice (P<0.05) while had little effect on intermediate monocytes. TLR9 inhibitor slightly decreased the percentages of the two subset monocytes but the effect did not reach statistical significance (**Fig. 3A**).

**Figure 3.**
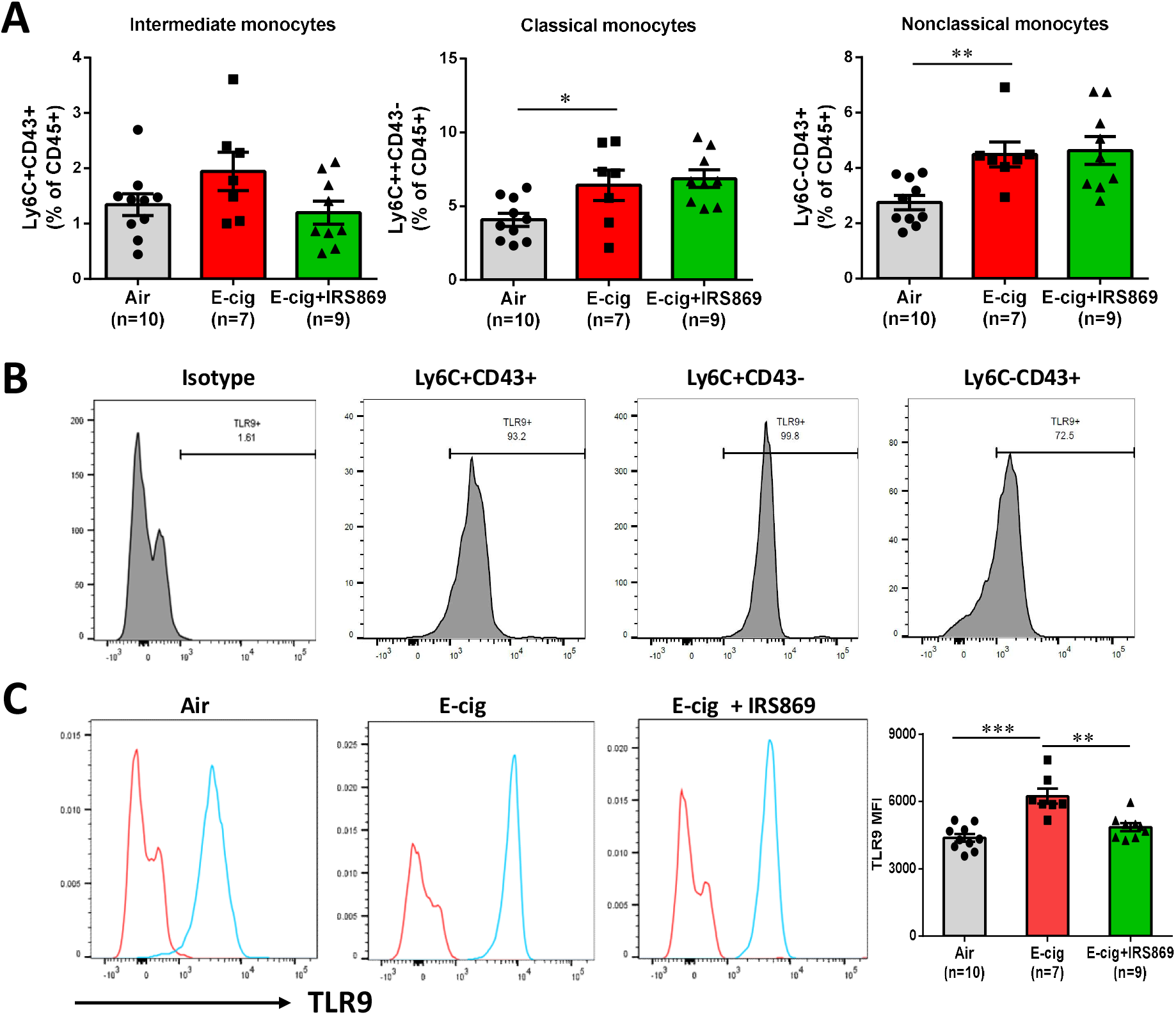
Effect of e-cig exposure on the percentages of different subsets of monocytes and TLR9 expression in the blood of ApoE^−/−^ mice. (A) Flow cytometric analysis of mouse blood cells. Singlet cells were firstly gated and then CD45+ cells were selected, followed with Ly6G+ neutrophil exclusion, CD11b+ and CD115+ cell selection. Finally, classical monocytes were gated as Ly6C+CD43-cells, nonclassical monocytes as Ly6C-CD43+ cells, and intermediate monocytes as Ly6C+CD43+ cells. (B) Representative histogram of TLR9 expression in different subsets of monocytes in air-exposed ApoE^−/−^ mice. (C) Mean fluorescence intensity (MFI) of TLR9 expression in classical monocytes. N means the number of mice in each group. Data are shown as mean ± SEM and statistical analysis was performed using the Students’ t-test. *P < 0.05; **P < 0.01; ***P < 0.001.

Activation of TLR9 has been reported to stimulate inflammatory responses of monocytes/macrophages and accelerate their transformation into foam cells, an indication of atherosclerotic plaque build-up.^14, 15, 37, 38^ We thus further examined TLR9 expression in the different subsets of blood monocytes. There was high frequency TLR9 expression in all three subsets of monocytes (87.6% ± 13.3%), with the lowest in nonclassical monocytes (79.7% ± 8.7%; **Fig. 3B**). Interestingly, classical monocytes from e-cig-exposed mice expressed very significantly (P < 0.001) higher intensity of TLR9 (mean fluorescence intensity (MFI), 6123.2 ± 1054.8) than cells from air-exposed mice (MFI 4415.8 ± 549.4; **Fig. 3C**). Pre-administration of mice with TLR9 antagonist IRS869 significantly reduced e-cig-upregulated TLR9 intensity (MFI 4974 ± 578.6; P<0.01) in classical monocytes. In contrast, there was lower frequency of TLR3 expression in all subsets of monocytes (27.4% ± 19.7%) and e-cig exposure had no significant effect on TLR3 expression (supplementary Fig. 2).

### TLR9 blockade decreased macrophage accumulation and the expression of adhesion molecule and TLR9 in atherosclerotic plaques

It has now been established that foam cells in atherosclerotic plaques are of bone marrow origin ^39^ although the mechanisms of monocyte recruitment into noninflamed aortas are not well defined. As shown in **Fig. 4A**, in addition to the upregulation of TLR9 expression in blood monocytes, ECV exposure significantly increased the expression of TLR9 in atherosclerotic plaques (P<0.05). The expression of adhesion molecule VCAM-1 was also significantly increased after e-cig exposure, both on the lesion surface and within the plaques (P<0.05, **Fig. 4B**). Pretreatment of mice with TLR9 inhibitor reduced the upregulation of both TLR9 and VCAM-1 in lesion sites. E-cig exposure also remarkably enhanced the accumulation of F4/80^+^ macrophages in atherosclerotic plaques, which could be decreased by TLR9 inhibitor administration (**Fig. 4C**).

**Figure 4.**
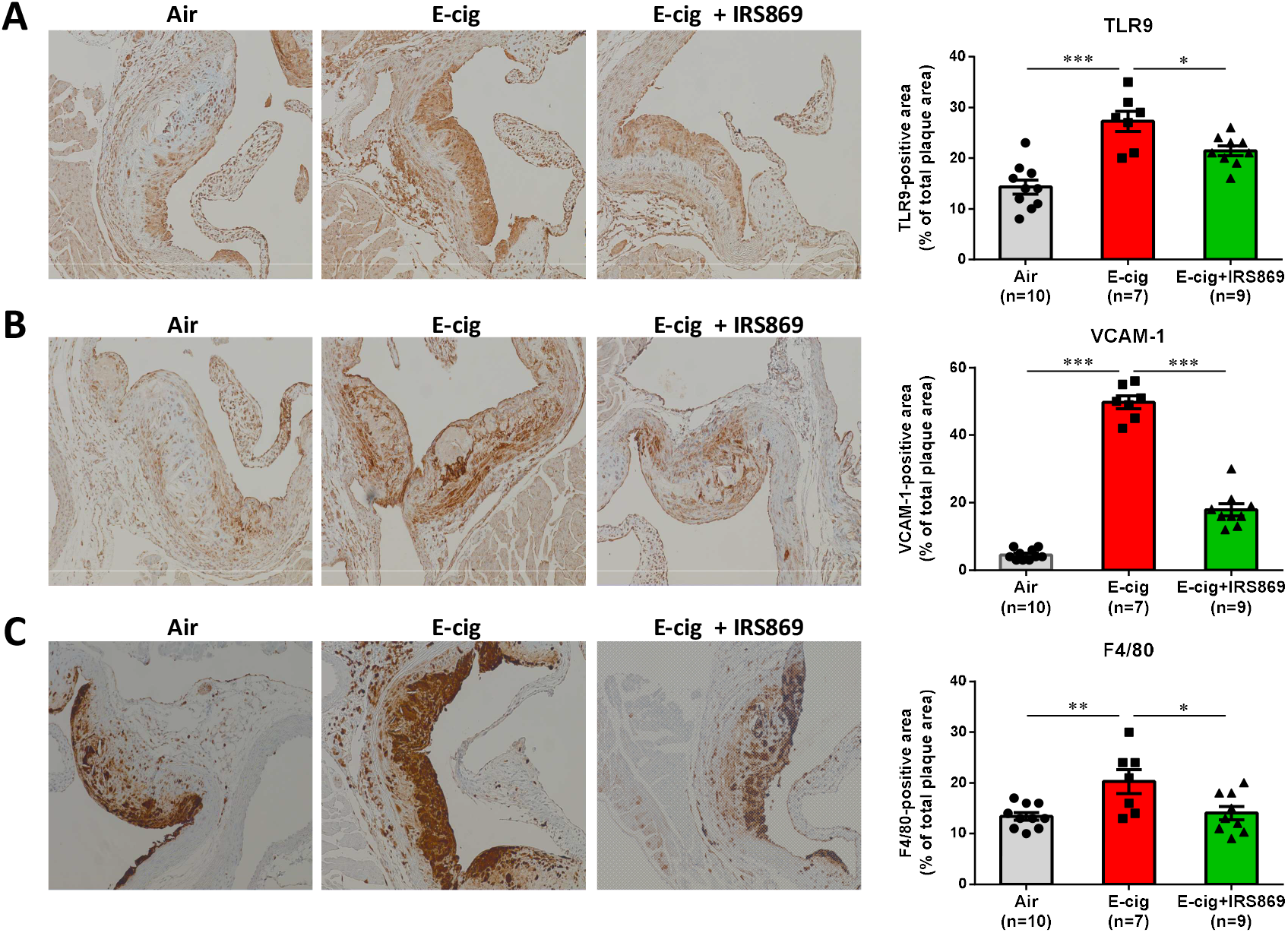
Pretreatment of mice with TLR9 inhibitor alleviated ECV exposure-induced macrophage accumulation and expression of adhesion molecule and TLR9 in atherosclerotic lesion. (A) Representative pictures of immunostaining of cross-sectioned aorta with TLR9 antibody (1:100) and quantification analysis between groups. (B) Representative pictures of immunostaining of cross-sectioned aorta with VCAM-1 antibody (1:100) and quantification analysis between groups. (C) Immunohistochemical staining of aortic cross-section with mouse macrophage marker F4/80 and macrophage content was expressed as percentage of F4/80-positive cells. N means the number of mice in each group. Data are shown as mean ± SEM and statistical analysis was performed using the Students’ t-test. *P < 0.05; **P < 0.01; ***P < 0.001.

### E-cig exposure increased CCR2^+^ cells in classical monocytes

Recent studies have suggested that Ly6C^high^ inflammatory monocytes are more readily to migrate into atherosclerosis-prone arteries through CCR2 or CX3CR1 chemokine receptors and then differentiate into aortic macrophages.^40^ It was found that e-cig exposure significantly increased the frequency of CCR2^+^ cells in classical monocytes (P<0.001) and pre-administration with TLR9 inhibitor could reduce the upregulation (P<0.05; **Fig. 5A**). On the contrary, the expression of CD62L in classical monocytes, which play an important role in lymphocyte-endothelial cell interactions was not affected either by e-cigs or the TLR9 inhibitor (P>0.05; **Fig. 5B**).

**Figure 5.**
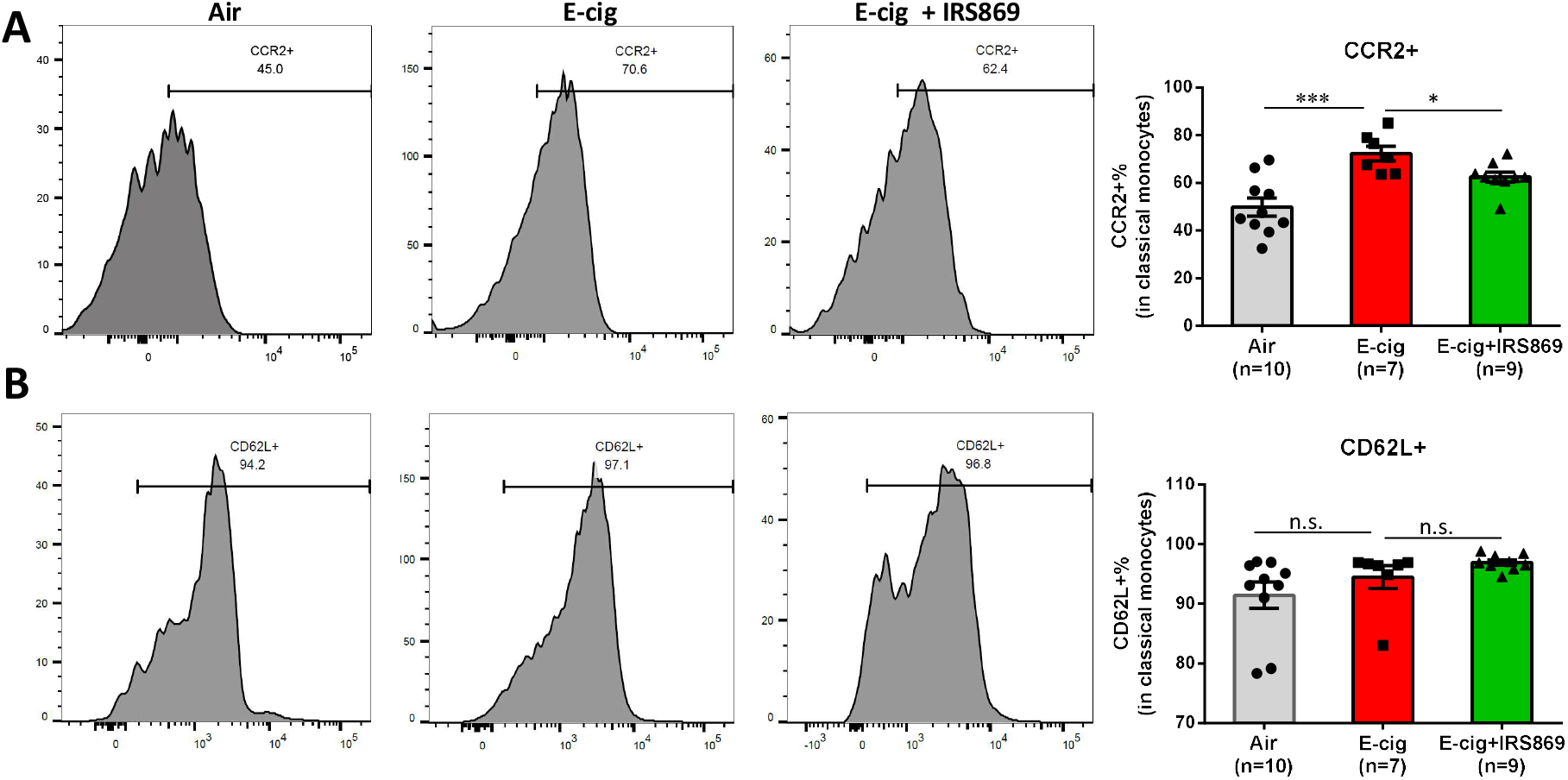
Effect of e-cig exposure on the frequency of classical monocytes expressing CCR2 and CD62L in ApoE^−/−^ mice. (A) Representative histograms of CCR2 expression in classical monocytes in air-exposed ApoE^−/−^ mice exposed to air, ecig, or e-cig with pre-administration of TLR9 inhibitor IRS869. (B) Expression CD62L expression in classical monocytes. N means the number of mice in each group. Data are shown as mean ± SEM and statistical analysis was performed using the Students’ *t*-test. *P < 0.05; ***P < 0.001; n.s.= not significant.

### ECV induces inflammatory response which can be suppressed by TLR9 inhibitor

The production of pro-inflammatory cytokines plays an important role in smoking-associated CVDs risks. **Fig. 6A-C** show that ECV exposure significantly increased the plasma levels of a number of proinflammatory cytokines, including TNF-a, RANTES, and IL-6 in ApoE^−/−^ mice (P<0.01). Pretreated the mice with TLR antagonist IRS 869 significantly suppressed e-cig-induced upregulation of these cytokines/chemokines in the plasma (P<0.05). Further study showed that pretreatment of murine monocyte-macrophages ex *vivo* with IRS869 could suppress the induction of cytokines by plasma from ECV-exposed mice (**Fig. 6D**). Together, these results suggesting the TLR9 activation is involved in plasma-induced monocyte activation and inflammatory response from ECV-exposed mice.

**Figure 6.**
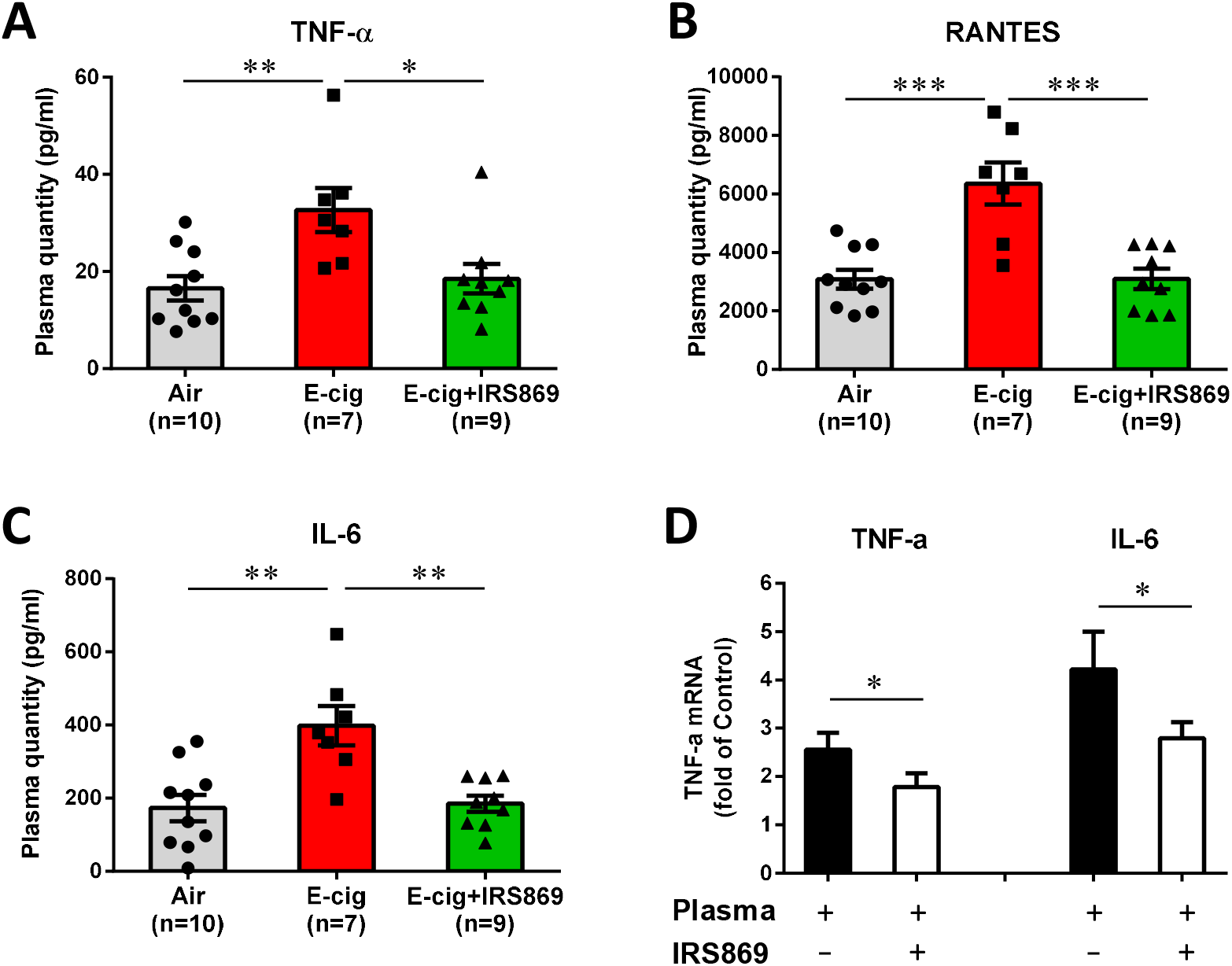
Effect of e-cig exposure on the plasma levels of inflammatory cytokines in ApoE^−/−^ mice and impact of plasma on cytokine expression in RWA264.7 cells *ex vivo*. (A, B, C) Plasma were collected from different groups of ApoE^−/−^ mice and subjected to immunoassay for a panel of inflammatory cytokines using the mouse Cytokine 9-Plex ELISA Kit. Images were scanned and the dot density was quantified by a parallel set of individual cytokine standards. N means the number of plasmas in each group used for cytokine measurement. Data are shown as mean ± SEM and statistical analysis was performed using the Students’ t-test. *P < 0.05; **P < 0.01; ***P < 0.001. (D) Effect of plasma from ECV-exposed mice on proinflammatory cytokines in murine monocytes/macrophages. RAW264.7 cells were pretreated with or without IRS869 at 5 μM for 30 min, then with plasma from ECV-exposed ApoE^−/−^ mice for 3 h. The expression of cytokines was examined by quantitative real time PCR. Data are shown as mean ± SEM and statistical analysis was performed using the Students’ t-test from three independent experiments using plasma from 3 mice. *P < 0.05.

### E-cig increased DNA damage of ApoE^−/−^ mice

TLR9 is an intracellular immune sensor of dsDNA, particularly oxidized DNA and GC-rich fragments than nonoxidized DNA fragments.^41^ Recently, e-cig aerosol was reported to damage DNA and impair the cellular antioxidant defenses.^30^ We thus examined the DNA damage in e-cig exposed mice with or without TLR9 inhibitor pretreatment. As shown in **Fig. 7A**, e-cig exposure significantly increased (2-fold) the plasma levels of 8-hydroxy-2’-deoxyguanosine (8-OHdG) in ApoE^−/−^ mice. 8-OHdG is one of the most representative products of oxidative modifications of DNA and has therefore been widely used as a biomarker for oxidative stress-related DNA damage. ECV exposure also greatly increased the mtDNA/nDNA ratio (1.59 ± 0.63 to 4.27 ± 1.49) in the plasma, indicating damaged DNA in the plasma mainly attributed to the degradation of oxidative damaged mtDNA (**Fig. 7B**). Intriguingly, unlike the ability to inhibit TLR9 activation and inflammatory response, pre-administration with TLR9 inhibitor could not reduce the DNA damage as compared to ECV exposure alone.

**Figure 7.**
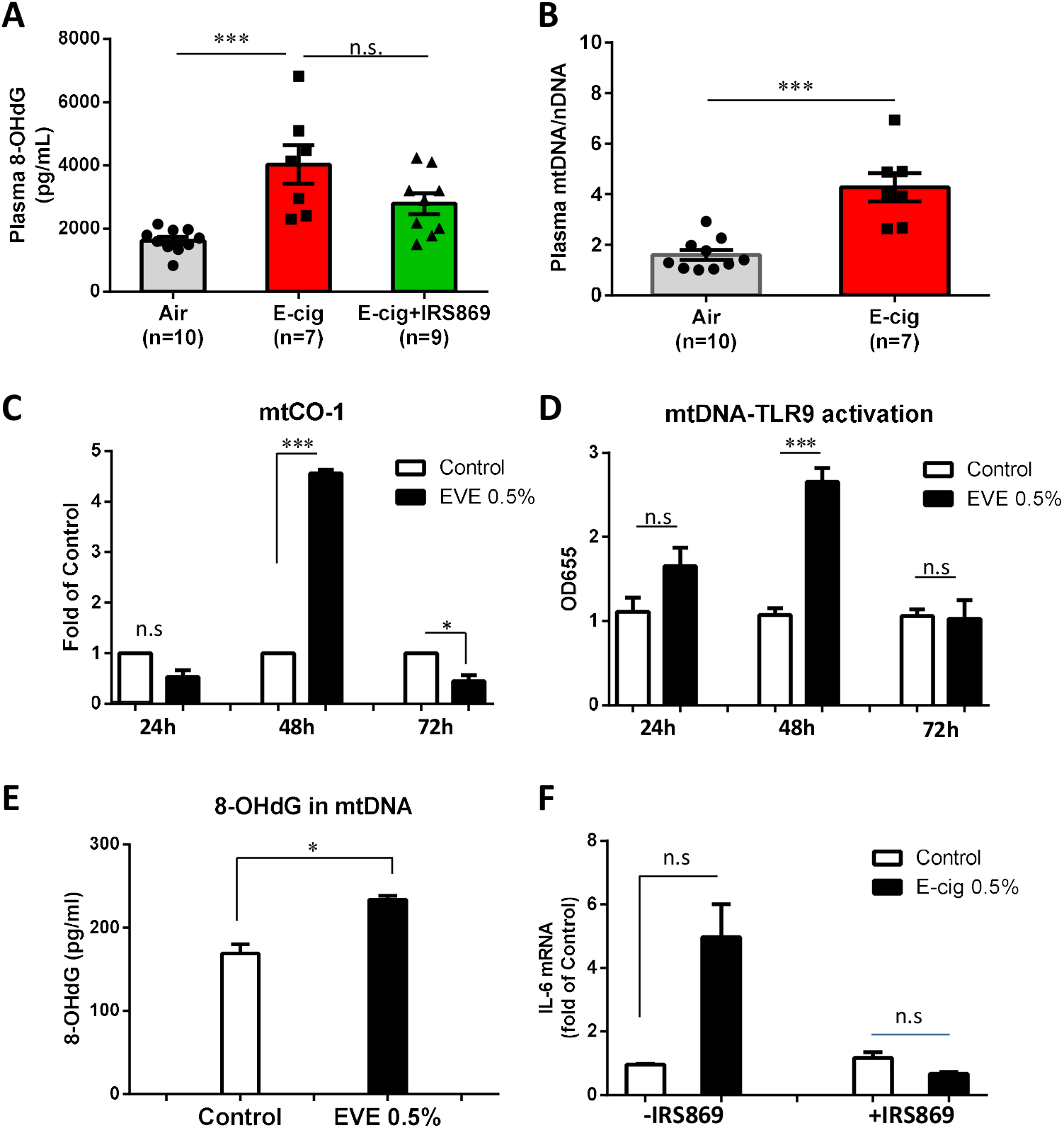
Effect of e-cig on mitochondrial DNA damage. (A) E-cig exposure increased plasma levels of circulating damaged DNA in ApoE^−/−^ mice. Plasma were collected from air-, e-cig aerosol-exposed ApoE^−/−^ mice with or without IRS869 preadministration. The plasma levels of circulating damaged DNA were measured by DNA damage ELISA kit and expressed by the plasma levels of 8-OHdG. (B) The plasma levels of circulating damaged DNA were measured by DNA damage ELISA kit and expressed by SYBR qPCR and the ratio of mtDNA against nDNA was calculated. N means the number of plasmas of each group used for DNA damage measurement. Data are shown as mean ± SEM and statistical analysis was performed using the Students’ t-test. ***P < 0.01. (C) ECV extract treatment upregulated cytoplasmic levels of mtDNA. RAW264.7 cells were treated with 0.5% of ECV extract for indicated period. The cytoplasmic mtDNA was extracted and the expression of mtCO-1 was measured by quantitative PCR. (D) HEK-Blue™ mTLR9 reporter cells were stimulated with 25 μg/ml of cytoplasmic mtDNA extracted from ECV extract-treated RAW264.7 cells after indicated time periods. After 24h incubation, NF-kB-induced SEAP activity was assessed using HEK-Blue™ Detection and reading the optical density (OD) at 655 nm. (E) The 8-OHdG levels in the cytoplasmic mtDNA was measured by Competitive DNA Damage ELISA. (F) Effect of pretreatment of macrophages with IRS869 on lesioned mtDNA-induced inflammatory cytokine expression. Data are shown as mean ± SEM from three independent experiments and statistical analysis was performed using the Students’ t-test. *P < 0.05; ***P < 0.001.

### Cytoplasmic mtDNA induced by ECV extract could activate TLR9

We next treated macrophages with ECV extract, extracted the cytoplasmic mtDNA and examined the functional significance of the damaged mtDNA in TLR9 activation. **Fig. 7C** shows that ECV extract treatment of RAW264.7 cells for 48 h increased the cytoplasmic mtDNA levels while there was no increase at 24 h and the levels diminished after 72 h. Correspondingly, cytoplasmic mtDNA from the 48 h ECV extract-treated macrophages significantly induced the activation of TLR9 in HEK mTLR9 reporter cells than that from control-treated RAW264.7 cells (**Fig. 7D**). To characterize the underlying mechanism of this difference, we examined the oxidative lesion in the cytoplasmic mtDNA. As shown in **Fig. 7E**, 8-OHdG level was significantly higher in cytoplasmic mtDNA from ECV extract-treated cells than that from control-treated cells, indicating that ECV extract treatment increased oxidative mtDNA lesion and the release of oxidized mtDNA into cytoplasma. Pharmacological inactivation of TLR9 could attenuate ECV extract-induced lesioned mtDNA-mediated inflammation in macrophages (**Fig. 7F**).

## Discussion

In the present study, our results show that ECV exposure induces atherosclerosis development, oxidative DNA damage and TLR9 activation in ApoE^−/−^ mice, substantiated by enhanced TLR9 expression in e-cig smoking-associated atherosclerotic plaques from femoral artery in e-cig smoker. In contrast, pharmacological inactivation of TLR9 can attenuate lesioned mtDNA-triggered TLR9 activation in macrophages and alleviate atherosclerosis *in vivo.*

In the past 20 years, immune and inflammatory mechanisms have gained tremendous interest in studying atherosclerosis.^1, 42, 43^ It is now well established that foam cells in atherosclerotic plaques are mainly derived from bone marrow^39^ and inflammatory monocytes are a major cellular component infiltrating into atherosclerotic plaque.^44–46^ It has been shown that increase of inflammatory monocyte subset (Ly6C^high^ monocytes) was associated with hypercholesterolemia in mice. As compared to Ly6C^low^ monocytes, Ly6C^high^ monocytes are more readily in migrating to atherosclerosis-susceptible arteries.^47^ We here demonstrated that e-cig-mediated atherosclerosis was accompanied with elevated frequency of both Ly6C^high^ classical monocytes and Ly6C^low^ nonclassical monocytes in blood and the accumulation of macrophages in plaques. The upregulation of monocytes was not affected by TLR9 inhibition although TLR9 inhibitor suppressed e-cig-mediated atherosclerosis development, suggesting there might be other cascades involved in mediating monocyte recruitment to endothelium. This hypothesis is supported by the finding that in TLR-mediated acute vascular inflammation, TLR9 agonist could not lead Ly6C^low^ monocytes to patrol the vascular wall.^48^ In addition, studies have suggested that Ly6C^high^ monocytes are not only prone to enter developing atherosclerotic lesions, but also differentiate into aortic macrophages via chemokine receptors, such as CX3CR1, CCR2, and CCR5.^40^ Our study shows the increased frequency of CCR2^+^ classical monocytes by ECV exposure could be restored by TLR9 inhibition. On the contrary, the expression of CD62L – a cell adhesion molecule that play an important role in lymphocyte-endothelial cell interactions^49^, was affected neither by e-cigs nor TLR9 inhibition. Thus, it is likely that in ECV-mediated atherosclerosis development, Ly6C^+^ monocytes utilized CCR2 to be recruited into inflamed arteries.

In addition to chemokine receptors that mediate monocyte trafficking, endothelial adhesion molecules (e.g. E- and P-selectin, and VCAM-1) also play important roles in the subendothelial accumulation of inflammatory monocytes.^50^ The upregulation of these molecules in lesion-prone areas is a hallmark of atherosclerosis.^51^ We here observed that e-cig-induced atherosclerosis was accompanied with upregulation of VCAM-1 expression in plaques. E-cig exposure also upregulated plasma levels of RANTES (CCL5) and several other inflammatory cytokines (TNF-α, IL-6). Our results support a previous notion that combined inhibition of CCL2, CX3CR1, and CCR5 could abrogate bone marrow monocytosis and almost abolish atherosclerosis in hypercholesterolemic, atherosclerosis-susceptible ApoE^−/−^ mice.^52^

Induction of DNA damage is a critical mechanism underlying smoking-associated tumor development, particularly in the pathogenesis of lung cancers.^22, 23, 25, 53^ Subjects with coronary artery disease (CAD) were found to have increased circulating nucleotide levels than healthy subjects.^54^ We observed that ECV exposure increased 8-OHdG levels in the plasma of ApoE^−/−^ mice. This result is consistent with previous studies showing that e-cig aerosols could suppress cellular antioxidant defenses or DNA-repair activity and induce significant oxidative DNA damage both *in vitro* and *in vivo*.^30, 31^ Our earlier study also showed that e-cig aerosols could dysregulate the oxidative phosphorylation (OXPHOS) complexes, resulting in the disruption of the nuclear/mitochondrial stoichiometry and mitochondrial dysfunction.^32^ In the present study, we further characterized that the increased circulating DNA in ApoE^−/−^ mice by ECV was enriched with mtDNA with oxidative DNA lesions compared to the genomic DNA. This results indicate that mtDNA is more susceptible to oxidative stress and consequently mtDNA is vulnerable to oxidation.^41, 55^

The correlation of mitochondrial oxidative stress (mitoOS) with the progression of human atherosclerosis has been reported. Excessive production of mitoOS was found in various cell types in atherosclerosis and with aging.^56^ Particularly, mitoOS in lesioned myeloid cells was found to enhance atherosclerosis development in aged mice.^57^ Cigarette smoke-induced mtDNA damage was proposed to promote atherogenesis in ApoE^−/−^ mice fed a chow diet, but underlying mechanism is not explored.^28^ Interestingly, in our study pre-administration with TLR9 inhibitor did not reduce plasma levels of 8-OHdG induced by e-cig, suggesting that inhibition of TLR9 activation but not oxidative stress-induced DNA damage underlies the suppression of inflammatory response and e-cig induced atherosclerosis development. As compared to nonoxidized DNA fragments, oxidized DNA and GC-rich fragments of mtDNA are stronger TLR9-stimulating ligands.^58^ Our *in vitro* study showed that ECV extract could increase the cytoplasmic mtDNA in murine RAW264.7 macrophages. The aberrant release of mtDNA into the cytosol is known as mtDNA stress and was reported with the ability to prime antiviral innate immune response through engaging DNA sensor cGAS.^59^ We found that the ECV extract-treated macrophages also exhibited mtDNA stress as evidenced by the increased levels of cytoplasmic mtDNA as compared to control-treated cells. More importantly, the cytoplasmic mtDNA from ECV extract-treated macrophages could more potently induce TLR9 activation than mtDNA from control cells. This TLR9 activation by e-cig-induced mtDNA stress is similar to a recent report showing that mitochondrial fission-induced mtDNA stress could activate TLR9 and initiate NF-κB signaling pathway, promoting tumor-associated macrophages (TAM) recruitment and polarization.^60^ Our further characterization of the mtDNA revealed that cytoplasmic mtDNA from ECV extract-treated cells had significantly higher levels of oxidative DNA marker 8-OHdG than that from control-treated cells. Due to the fact that oxidized DNA is stronger TLR9 stimulus than nonoxidized DNA,^58^ the higher potency to activate TLR9 by mtDNA from ECV extract-treated cells implicates that not only mtDNA stress but also oxidative mtDNA lesion underlie e-cig-triggered TLR9 activation and atherosclerosis development in ApoE^−/−^ mice.

The pivotal role of TLR9 has recently been reported in angiotensin II-induced atherosclerosis.^18^ There was increased circulating levels of cfDNA in angiotensin II – infused ApoE^−/−^ mice whereas TLR9 blockade could suppress the proinflammatory activation of macrophages and vascular inflammation.^61^ In addition, the activation of TLR9 has been indicated by hepatocyte mtDNA in driving nonalcoholic steatohepatitis.^62^ More relevantly, SHR were found to have elevated circulating mtDNA, which can activate TLR9 and contribute to vascular dysfunction.^21^ TLR9 activation of RAW264.7 cells was shown to stimulate the inflammatory responses and accelerate the transformation into foam cells.^63^ High Dose CpG ODN1826, an agonist ofTLR9, increases atherosclerotic plaque formation^16^ and pharmacological TLR9 inactivation can alleviate atherosclerotic progression^17^, suggesting a pro-atherosclerotic role of TLR9. Despite the above reports on the pro-atherosclerotic role of TLR9, an atheroprotective function of TLR9 is implicated by the conventional genetic deletion of TLR9 exacerbates atherosclerosis in ApoE^−/−^ mice with high-fat diet by a 33% increase in lipid deposition and plaque size, and administration of TLR9 agonist reduced lesion severity.^64^ These controversial results implicate that the atherogenic effect of intracellular TLRs, including TLR9, might be context or cell type-dependent. In 2013, Lundberg et al. showed LDLR^−/−^:TLR3^−/−^ bone marrow (BM) chimeric mice were protected from atherosclerotic lesion formation when fed with high-fat diet. This result suggests that selectively TLR3-deficient in hematopoietic cells is atheroprotective rather than atheroprone in wholebody knockout situation.^65^ It also supports another report showing that selective deficiency of IFN-β in BM decreases atherosclerotic lesion formation.^66^ These results together implicate that hematopoietic intracellular TLR signaling is detrimental to atherosclerotic lesion development.^67, 68^ We here showed that e-cig significantly enhanced TLR9 intensity in circulating monocytes and TLR9 expression in atherosclerotic plaque, and administration of a specific TLR9 antagonist IRS869 prior to e-cig exposure could significantly reduce the atherosclerotic lesions in ApoE^−/−^ mice, demonstrating a pro-atherosclerotic role of TLR9 in this setting.

In conclusion, our study shows that oxidative mtDNA damage and TLR9 activation of monocytes are critical mechanisms involved in e-cig-accelerated atherosclerosis development in ApoE^−/−^ mice. Intervention of TLR9 activation is a potential treatment of e-cig aerosol-related inflammation and CVDs.

## Author Contributions

Conceived and designed the experiments: JL, HW. Performed the experiments: HLD, HJ, HS, WC, TLTL. Analyzed the data: JL, HLD, HW. Contributed reagents/materials/analysis tools: BK, TR, MS, KP, CL, MST. Wrote the paper: JL, MST, HW.

## Acknowledgments

We acknowledge Dr. Michael Autieri from Temple University in providing the murine macrophage RAW264.7 cells. Dr. Gratian Salaru from the Flow Cytometry Core of Robert Wood Johnson Medical School of Rutgers University is acknowledged for flow cytometry data analyses. Financial support from Bristol-Myers Squibb (New Jersey) in establishing the Temple Heart Transplantation Biorepository is also acknowledged. This study is partly supported by Cancer Center Support Grant P30CA072720.

