## Supplemental Materials for "Electronic cigarettes induce mitochondrial DNA damage and trigger toll-like receptor 9-mediated atherosclerosis"

Short title: MtDNA-TLR9 mediates e-cig-induced atherosclerosis

Jieliang Li[1,#], Do Luong Huynh[1,#], Moon-Shong Tang[2], Hannah Simborio[3], Jing Huang[1], Beata Kosmider[3,4], Michael B. Steinberg[5], Le Thu Thi Le[1], Kien Pham[6], Chen Liu[6], He Wang[1,*]

***Correspondence to:** He Wang, MD, PhD, 675 Hoes Ln, Department of Pathology and Laboratory Medicine, Robert Wood Johnson Medical School, Rutgers University, Piscataway, NJ 08854 USA Tel: 732-937-8651; Fax: 732-507-8115

**This PDF file includes:**

Supplementary text
Figures S1 to S2
Tables S1

### Supplementary Information Text

### Materials and Methods

#### Animals

Animal handling and experimentation were in accordance with the recommendation of the current NIH guidelines and were approved by the Temple University School of Medicine Institutional Animal Care and Use Committee. C57BL/6J A*poE*$^{-/-}$ mice of both sexes were purchased from the Jackson Laboratory (Bar Harbor, ME) and were housed in an inhalation facility under controlled conditions of temperature, humidity, and a pathogen-free environment in the animal facility of Temple University School of Medicine for one week acclimatization before whole body exposures. All the mice were maintained under a 12:12 light:dark cycle (lights on between 6 AM to 6 PM). The animals were kept in cages and were fed a regular diet (pelleted food) and water *ad libitum*, unless otherwise indicated. The animals were monitored on a daily basis during exposure to check for any signs of distress.

#### Measurements of cotinine levels

Urine cotinine levels were monitored at two time points during the 16-week experiment, one after the 3rd day exposure and one at the endpoint. Briefly, within 1 h after exposure, mouse urine was collected and used for determination of cotinine levels using the Calbiotech Cotinine ELISA kit (El Cajon, CA) according to manufacturer’s instructions.

#### Cholesterol and lipoprotein measurement in serum

Aliquots of plasma were analyzed using commercial kits from Thermo Clinical Labsystems (Frankfurt, Germany) according to manufacturer's instructions.

### Flow cytometry

Whole blood in a 1.5 mL Eppendorf tube was centrifuged for 10 min at 300 ×*g* and at room temperature. Plasma was then removed and stored at -80ºC for future analysis. The cell pellet was washed once with FACS buffer and then mixed with 50 µL of ice cold FACS buffer containing mouse Fc Block (BD Biosciences, San Jose, CA) and incubated at room temperature for 30 min. After centrifugation and discarding the supernatant, cells were mixed with 100 µL antibody cocktail containing the following fluorescently conjugated antibodies for cell surface markers: BUV395-conjugated anti-CD3 MAb 17A2 (Cat#563565), BUV395-conjugated anti-CD19 MAb 1D3 (Cat#563557), BUV395-conjugated anti-NK1.1 MAb PK136 (Cat#564144), BB515-conjugated anti-MHCII MAb IE/IA 2G9 (Cat#565254), BV421-conjugated anti-CD45 MAb 30-F11 (Cat#563890), BV510-conjugated anti-CD62L MAb MEL-14 (Cat#563117), BV605-conjugated anti-CCR5 C34-3448 (Cat#74369), phycoerythrin (PE)-Cy7-conjugated anti-CD43 MAb S7 (Cat#562866), allophycocyanin (APC)-Cy7-conjugated anti-Ly6C MAb AL-21 (Cat#560596) from BD Biosciences, Alexa700-conjugated anti-CCR2 MAb 475301 (FAB5538N-100), Alexa647-conjugated anti-TLR9 MAb 1138D (FAB7960R-100 µg) from R&D Systems (Minneapolis, MN), Peridinin-Chlorophyll-Protein (PerCP)-Cy5.5-conjugated anti-CD115 MAb AFS98 (Cat#135525), PE-conjugated anti-TLR3 MAb 11F8 (Cat#141903) from BioLegend (San Diego, CA). Isotype controls used were BB515-conjugated rat IgG2a (Cat# 564418), BV421-conjugated rat IgG2b (Cat# 562603), BV510-conjugated rat IgG2a (Cat# 562952), PE-Cy™7-conjugated rat IgG2a (Cat# 552784), APC-Cy™7-conjugated rat IgM (Cat# 560571) from BD Biosciences, BV605-conjugated rat IgG2c (Cat#400727), PE-conjugated rat IgG2a (Cat# 400507), and PerCP/Cy5.5 rat IgG2a (Cat#400532) from BioLegend, Alexa647-conjugated rabbit IgG (Cat#IC1051R), and Alexa700-conjugated rat IgG2b (Cat#IC013N) from R&D Systems. Detailed sources and controls Cells were incubated for 30 min on ice in the dark. After centrifugation, the cell pellets were mixed with ice cold ACK lysis buffer (ThermoFisher) and incubate at room temperature for 10 min and in the dark. This will lyse the RBCs and simultaneously wash the cells from the staining. Finally, cells were fixed in 2% paraformaldehyde in ice cold FACS buffer for 10 min, washed, and then resuspended in 300 µL of FACS buffer. Analysis of stained cells was performed with a FACSCanto LSRII flow cytometer (BD Biosciences). Isotype controls were used to set appropriate gates. Data were analyzed acquired with FACSDiva (BD Biosciences) and analyzed with FlowJo 6.4.7v10.6 (Tree Star, Ashland, OR). For all samples, approximately 20,000 cells were analyzed to generate scatter plots. All events were collected from each sample to generate the scatter plots.

### *Ex vivo* treatment of macrophages with plasma, RNA extraction and quantitative real time PCR

Murine monocyte/macrophage RAW264.7 cells (a gift from Dr. Michael Autieri, Temple University) were cultured in DMEM basic medium (Gibco, USA) supplemented with 10% fetal bovine serum (Gibco), 100 IU/mL penicillin and 100 µg/mL streptomycin at 37°C in a humidified atmosphere with 5% $CO_2$. RAW264.7 cells were pretreated with IRS869 (5 µM) for 1 h before further treatment with 10% (v/v) of plasma from e-cig-exposed ApoE$^{-/-}$ mice. After treatment, total RNA was extracted using the RNAqueous®-Micro Kit (Ambion, Austin, TX) and quantitated using a nanodrop spectrophotometer (Thermo Scientific, Waltham, MA) and was followed immediately by cDNA synthesis using the First Strand cDNA synthesis kit (Qiagen, Germantown, MD).

Quantitative real time PCR for genes of interest was performed using SYBR Green Master mix (Applied Biosystem, CA). Amplification conditions were: 95°C for 3 min, 95°C for 15 s, 55°C for 15 s, and 72°C for 30 s for 40 cycles. The CT value for glyceraldehyde-3-phosphate dehydrogenase (GAPDH) was subtracted from the CT value for the gene of interest to obtain a delta CT (ΔCT) value. The relative fold-change for each gene was calculated using the $2^{-\Delta\Delta Ct}$ method, and a melting-curve analysis was performed to ensure the specificity of the products. The primers used were shown in Supplemental Table 1 and synthesized by IDT.

**Preparation of ECV extract**

The ECV extract was prepared with a slight modification as previously described to prepare aqueous extract from conventional cigarettes (70). Briefly, e-cig aerosol was generated by V2 E-liquid (30.3 µL, containing ~0.72 mg nicotine, comparable to nicotine from the smoke of one 3R4F cigarette without filter (Kentucky Tobacco Research & Development Center, Lexington, KY) in a Kangertech EVOD Mega pen operated at a fixed voltage of 3.7V. The aerosol was drawn into 12.5 mL DMEM with peristaltic pump (Manostat 72-310-000; Barnant Company, Barrington, IL). The pump was set at an optimum speed to allow the E-liquid to burn in approximately 15 min and resulting solution was considered 100% ECV extract. This solution was filtered through a 0.22 µm pore acrodisc syringe filter and aliquot into 1 mL/tube and stored at -80°C. Nicotine concentrations from each batch of preparation were measured at the lab of Matthew S. Halquist (Bioanalytical Shared Resource Laboratory, University School of Pharmacy).

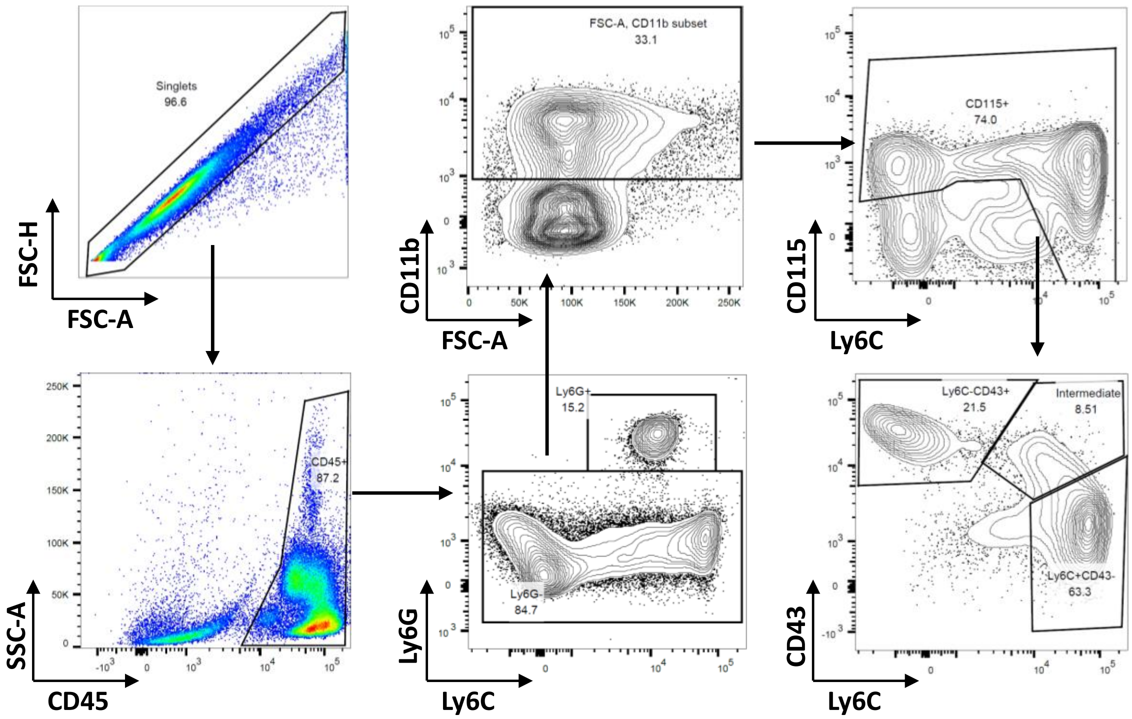


**Fig. S1. Flow cytometric analysis of mouse blood cells.** Singlet cells were firstly gated and then CD45+ cells were selected, followed with Ly6G+ neutrophil exclusion, CD11b+ and CD115+ cell selection. Finally, classical monocytes were gated as Ly6C++CD43- cells, nonclassical monocytes as Ly6C-CD43+ cells, and intermediate monocytes as Ly6C+CD43+ cells.

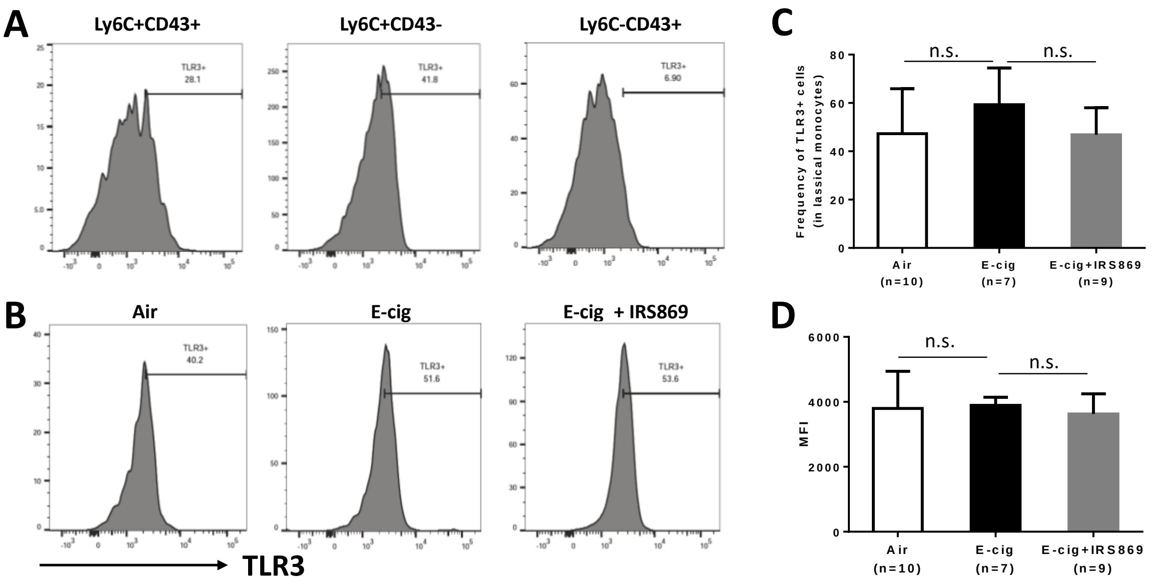


**Fig. S2. Effect of E-cig exposure on the expression of TLR3 at both frequency and intensity levels in blood monocytes in ApoE$^{-/-}$ mice. A.** Representative histogram of TLR3 expression in different subsets of monocytes in air-exposed ApoE$^{-/-}$ mice. **B, C.** Representative histograms of TLR3 expressing frequency in classical monocytes in different treatment groups of mice (B) and statistical analysis (C). **D.** Mean fluorescence intensity (MFI) of TLR3 expression in classical monocytes. Data are shown as mean ± SEM. n.s.=not significant.

**Table S1. Primers used in this study**

| Gene name | | Sequence (5'-3') |
|---|---|---|
| mCCL2 | F | GAAGCTGTGATCTTCAGGAC |
| | R | GGAACTCTCAGCCTAGACA |
| mCCL5 | F | CTCACCATATGGCTCGGA |
| | R | CGAGTGACAAACACGACTG |
| mIL-1 β | F | CTTTGAAGAAGAGCCCATCC |
| | R | TTCATCTCGGAGCCTGTAG |
| mIL-6 | F | GAGGATACCACTCCCAACAGACC |
| | R | AAGTGCATCATCGTTGTTCATACA |
| mIL-8 | F | CTCTTGGCAGCCTTCCTGATT |
| | R | TATGCACTGACATCTAAGTTCTTTAGCA |
| mIL-10 | F | TGAGGCGCTGTCGTCATCGATTTCTCCC |
| | R | ACCTGCTCCACTGCCTTGCT |
| mIL-12α | F | CTGCACTGCTGAAGACATC |
| | R | CTCCCTCTTGTTGTGGAAG |
| mIL-12β | F | GGGACATCATCAAACCAGAC |
| | R | TGAGGGAGAAGTAGGAATGG |
| mTNF-α | F | CACGTCGTAGCAAACCACCAAGTGGA |
| | R | TGGGAGTAGACAAGGTACAACCC |
| mGAPDH | F | GTCGGTGTGAACGGATTTG |
| | R | GTGAGTGGAGTCATACTGGA |
| m18S rDNA | F | TAGAGGGACAAGTGGCGTTC |
| | R | CGCTGAGCCAGTCAGTGT |
| mtCO-1 | F | GCCCCAGATATAGCATTCCC |
| | R | GTTCATCCTGTTCCTGCTCC |
| mTLR9 | F | GCTGTCAATGGCTCTCAGTTCC |
| | R | CCTGCAACTGTGGTAGCTCAC |
